# Grid cells anticipate the animal’s future movement

**DOI:** 10.1101/2024.12.05.627046

**Authors:** Dóra Éva Csordás, Johannes Nagele, Martin Stemmler, Andreas V. M. Herz

## Abstract

Grid cells in the rodent medial entorhinal cortex preferentially fire spikes when the animal is within certain regions of space. When experimental data are averaged over time, spatial firing fields become apparent. If these firing fields represented only the current position of the animal, a grid cell’s firing should not depend on whether the animal is running into or out of a firing field. Yet many grid cells are sensitive to the animal’s direction of motion relative to the firing-field center. Such apparent egocentric "inbound-outbound tuning" could be a sign of prospective encoding of future position, but it is unclear whether grid cells code ahead in space or in time. To investigate this question, we decided to undo the inbound-outbound modulation by shifting all spikes within a given firing field by a fixed distance in space or in time. For grid-cell data recorded in mice, optimizing in space requires a forward shift of around 2.5 cm, whereas optimizing in time yielded a forward shift of about 170 ms. In either case the firing-field sizes decrease. Minimizing just the field size yields somewhat smaller shifts (roughly 1.8 cm and around 115 ms ahead). Jointly optimizing along the temporal and spatial dimension reveals a continuum of flat inbound-outbound tuning curves and a shallow minimum for field sizes, located at about 2.3 cm and 35 ms. These findings call into question a purely spatial or purely temporal interpretation of grid-cell firing fields and inbound-outbound tuning.

## Introduction

Navigating through space requires a well-organized network of spatially active cells, such as place cells in the hippocampus (O’Keefe et al., 1971), head-direction cells in the postsubiculum (Taube et al., 1990), grid cells in the medial entorhinal cortex (Hafting et al., 2005), boundary vector cells in the subiculum (Lever et al., 2009) and speed cells in the medial entorhinal cortex (Kropff et al., 2015).

The traditional working hypothesis that these cells encode the current values of the dynamic variables they are tuned to, was already shown to be incorrect. Head direction cells, for example, anticipate future head directions by up to 95 ms (Stackman and Taube, 1998), and place and grid cells anticipate future spatial locations by 30-120 ms (Muller and Kubie, 1989, Sharp 1999) and 50-80 ms (Kropff et al., 2015), respectively. In addition, the cells’ coding properties, as inferred from the experimental data, may reflect implicit assumptions about the neural code. In fact, Huxter et al. (2008) showed that shifting the measured spike positions forward along the movement direction can shrink the calculated place field size. As a consequence, the calculated anticipatory activity of spatially tuned cells is influenced by the experimentalist’s choice about where to place the tracking LEDs on the animal’s head – the chosen LED position may or may not reflect the "true" position perceived by the animal.

To disentangle spatial effects from temporal anticipation, considering velocity-dependencies might help. This approach has proven successful in other studies and showed that head-direction cells in the anterodorsal nucleus and in the lateral mammillary nuclei systematically shift their directional firing preference as a function of angular velocity (Blair and Sharp, 1995, Stackman and Taube, 1998). Place cell firing correlates with future locations at slow speeds, but at high speeds it correlates with past locations (Sharp, 1999). Studying anticipation in animals foraging in 2D or 3D comes with an added benefit compared to 1D movements in that spatial and temporal aspects decouple as soon as the animal moves on a curved path.

It has remained an open question whether place cells or grid cells indeed show true anticipation or whether the observed phenomena are just a side effect of incorrectly assigning the animal’s perceived position on the body. Focusing on the medial entorhinal cortex, the present study explores whether the spatial position shifts, temporal anticipation or possibly a combination of both scenarios best explain grid cell firing.

## Materials and Methods

### Data

We analyzed data reported in Latuske et al. (2015). The dataset contained tetrode data (Sampling rate: 20 or 24 kHz) obtained from male mice during movements in a square arena (70 x 70 cm). The trajectory was recorded by three LEDs placed on the head of the animal, two at the ears, and one centrally positioned close to the animal’s neck. The spatial position of the animal was defined as the midpoint between the two LEDs at the ears (as shown in Fig. 1A). The head direction was determined by the vector pointing from the neck LED to this midpoint.

**Figure 1.**
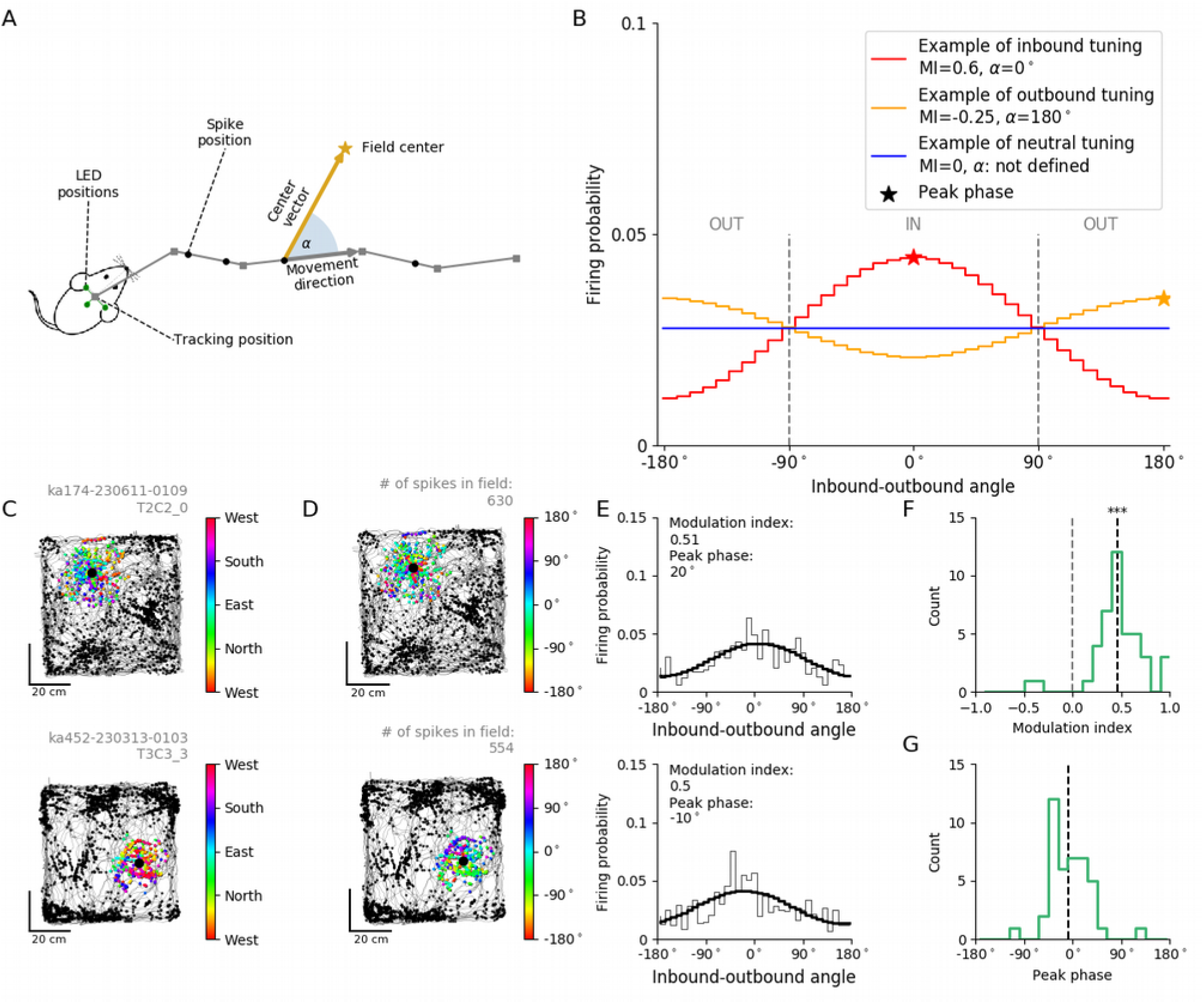
Inbound-tuned firing pattern of mouse grid cells A, Computation of the inbound-outbound angle a. There were three LEDs (green dots) placed on the head of the animal, from which the tracking position (grey squares) was computed at every sampling point. The positions of the spikes generated by a cell (black dots) were calculated from its spike times by interpolation along the animal trajectory. For all spikes of a particular grid field, inbound-outbound angles were computed as the angles between the momentary movement direction and the vector pointing from the spike position to the center of that grid field. B, To characterize the inbound-outbound tuning curves, modulation indices (MIs) and peak phases were calculated. For illustrative purposes, three qualitatively different fits are shown, inbound tuned (red), outbound tuned (orange) and a neutral case. The MIs show the tuning strength and their signs tell whether the tuning is inbound (+) or outbound (-). Asterisks denote the fits’ peak phase. C, Absolute running directions of an animal when spikes were fired for two example grid fields. (Grey: trajectory, black: spikes that are not members of the grid field, large black dot: center of the grid field, colored dots: spikes that are members of the firing field, colored by the absolute running direction). D, Inbound-outbound, egocentric running direction of an animal when spikes were fired. Same two example grid fields as in C (Spikes belonging to the particular grid field are colored by the running direction relative to the grid field center (inbound-outbound angle). E, Inbound-outbound firing probabilities for the two example grid fields, together with their cosine fits. (Thin black line: binned firing probability, thick black line: cosine fit, black text: MI and peak phase of the cosine fit). F, Relative frequency of modulation indices of the significantly tuned grid fields. (Grey dashed line: expected mean of the modulation indices, i.e. zero, black dashed line: Mean of the modulation indices. 3 asterisks note that the mean of the distribution of MI-s is significantly different from 0, T-test, p<0.01). G, Relative frequency of peak phases of the significantly tuned grid fields. (Black dashed line: Circular mean of the peak phases).

### Grid cell selection

After removing recording sessions for which the animal trajectories showed artifacts, 522 principal cells were identified using the same criteria as in Latuske et al. (2015); in particular, the mean firing rate had to be smaller than 10 Hz. Out of those cells, 115 cells had been classified as grid cells by Latuske et al..

### Grid field detection

The spikes belonging to each grid field ("member spikes") and the associated grid-field centers were assigned using the Mean Shift algorithm (Comaniciu and Meer, 2002). The Mean Shift algorithm and the estimation of its bandwidth (number of samples = 10) was performed using the Python implementation from the sklearn.cluster package (using default parameters otherwise). To account for temporal effects, each spike data was weighted by the dwell time of the animal in the corresponding spatial bin (2x2 cm) before clustering. The radius of a grid field was computed as the maximal distance of the member spikes from the respective grid field center. To avoid boundary effects, we focused on well-defined grid fields away from the arena’s boundary. To this end, grid fields were only considered further if the disk drawn around the grid-field center with the field radius was fully within the arena. Based on these criteria, 45 grid cells with 55 grid fields were included in the study (for 36 grid cells with 44 grid fields, the head direction was also recorded).

### Inbound-outbound firing probability in a grid field

To compute inbound-outbound firing probabilities, every spike in a given grid field was assigned an angle α which was the angle between the momentary velocity vector and the vector pointing from the spike position to the grid field center (see Fig. 1A). With this convention, 0 ° means that the animal was heading directly towards the grid field center when the spike was generated and +/-180° means that the grid field center was straight behind the animal. Furthermore, a positive angle means that the grid field center was on the left side of the animal and a negative angle means that it was on the animal’s right side. Accordingly, if α was between -90° and 90° the spike was fired on a trajectory segment on which the animal moved closer to the firing-field center and which will be called an "inbound trajectory", otherwise, the animal moved on an "outbound trajectory". The trajectory points themselves were assigned an angle in the same way. Polar histograms with 36 ten-degree bins in the range of -180° to 180° of both the spikes and the trajectory points were computed. The inbound-outbound firing rate was defined as follows:

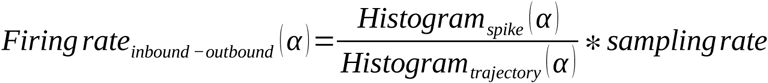

To correct for different firing rates in the grid fields, firing probabilities, whose sum over angular bins equals unity, were calculated by normalizing the firing rates.

### Inbound-outbound tuning in a grid field

The firing probability curves were fit by the least squares method with the following cosine model:

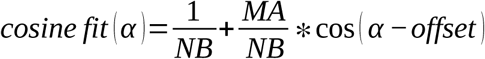

where NB was the number of bins (36) and the free parameters were the angular offset and the modulation amplitude (MA), which was assumed to be positive or zero. From the MA and offset we computed two related measures, i.e. the modulation index (MI) and the peak phase. If the peak of the cosine fit was between -90° and 90° the MI was defined to be equal to the absolute value of MA, otherwise MI was set to minus the absolute value of MA. The peak phase was the center of the bin at which the cosine fit had its maximum value. With these definitions, both the MI and the peak phase carry information about a grid cell was inbound or outbound tuned for the grid field under consideration (Fig. 1B).

### Statistical validation of a grid field being inbound-outbound tuned

To decide whether a grid field is significantly tuned we shuffled the bins of the original firing probability curve of the grid field 10000 times and fitted each shuffle again with the cosine model. If the grid field’s MI was smaller than the 5^th^ percentile or higher than the 95^th^ of the shuffled MI distribution, then the field was called significantly tuned. 42 out of 55 grid fields were significantly tuned. If the peak phase was less than +/-45° away from 0° or 180°, then the tuning curve was called centered. We found 30 grid fields with significantly tuned and centered tuning curves.

### Modeling different prospective or retrospective scenarios

The following scenarios were applied at the level of individual grid cells.

#### Movement direction (MD)

We shifted every trajectory point backwards (negative shift) or forwards (positive shift) along the animal’s momentary movement direction (MD). The MD was computed as the orientation of the average of the vectors defined by the previous and the following trajectory segments (the vector pointing from (px_i-1_,py_i-1_) to (px_i_,py_i_) and the vector pointing from (px_i_,py_i_) to (px_i+1_,py_i+1_)), where i denotes the i’th location in the raw position data set. Using the original spike times we recomputed the MD-shifted spike positions by interpolation on the MD-shifted trajectory (Fig. 2A first panel). The explored shifts ranged from -6 to +6 cm in steps of 0.2 cm.

**Figure 2.**
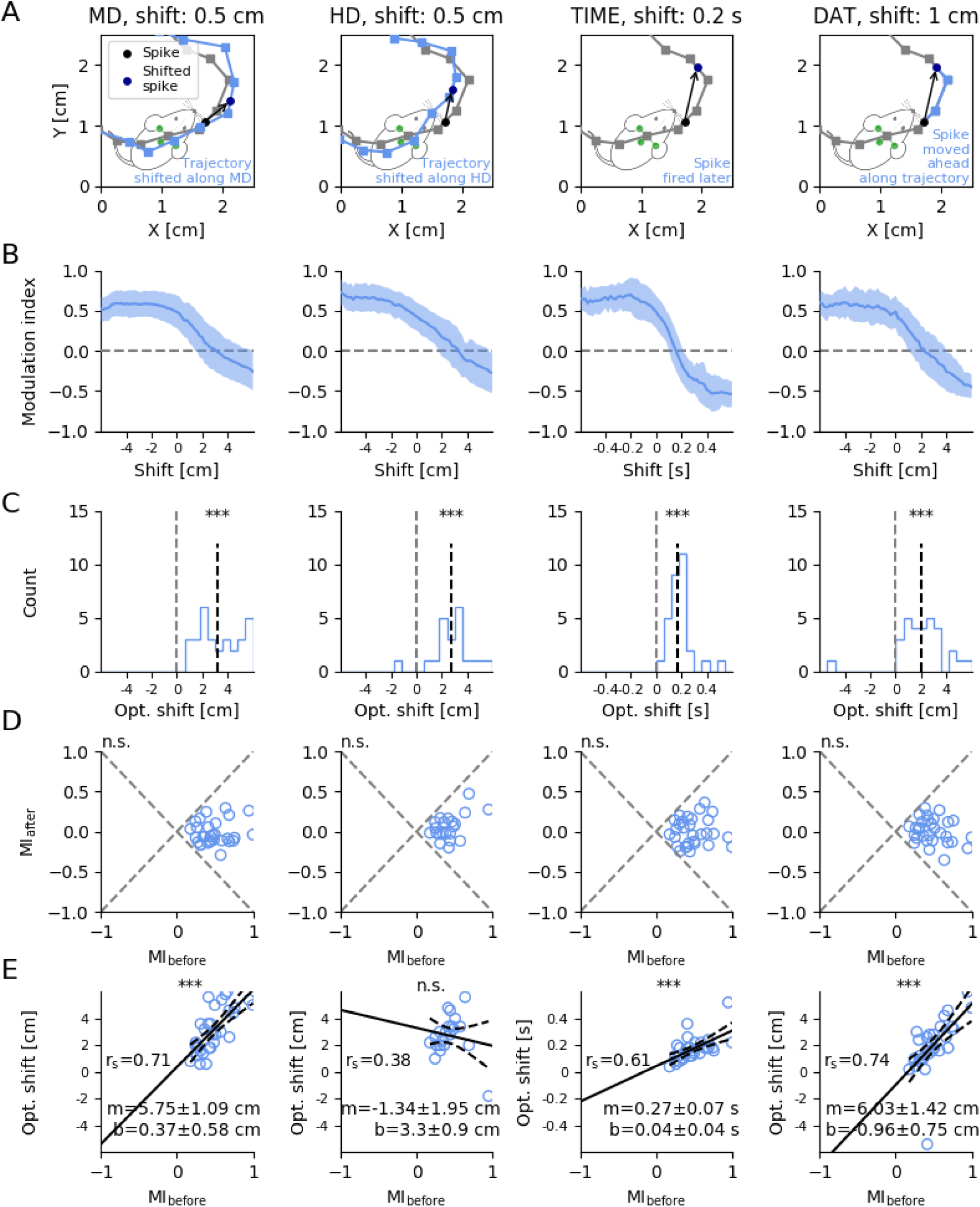
Alternative scenarios to flatten inbound-tuned firing rates A, Visualization of the four different scenarios that were used for optimization. For MD and HD we shifted the trajectory points forwards and backwards along the movement direction or head direction and kept the spike times. For TIME we assumed that spikes were fired with a positive of negative temporal offset, so we shifted the spike times and remapped them on the original trajectory. For DAT we moved spikes by a given distance along the trajectory and recomputed the spikes times from the new spike positions. B, Mean optimization curves in the different scenarios of the significant and centered fields. MI is shown as a function of the tested shift (Blue line: mean MI at shift, shaded area: SD around the mean, Grey dashed line: optimal modulation index). C, Distributions of optimal shifts in the different optimization scenarios. (Black dashed line: Mean of optimal shifts; three asterisks indicate that the mean of the distribution is highly significantly different from zero, T-test p<0.01/4) D, Modulation indices before and after optimization in the different optimization scenarios. The modulation indices are decreased after optimization and their means where never significantly different from zero (n.s.: not significant, T-test, p>0.05/4) (Grey dashed lines: lines of constant performance in terms of absolute MI values.) E, Optimal shifts as a function of MIs before shifting. The fit of regression lines were highly significant for the reliability of fit for MD, TIME and DAT. Black solid line: regression line, black dashed line: 95% confidence interval, r_s_: Spearman correlation value, m: slope of the regression line, b: intercept of the regression line, SD: standard error of the estimate. Three asterisks indicate that the Spearman correlation is highly significant (p<0.01/4) and n.s. notes not significant (p>0.05/4).

#### Head direction (HD)

We shifted every trajectory point along the animal’s head direction (HD), given by the vector pointing from the neck LED to the midpoint between the two LEDs at the ears. Using the original spike times we recomputed the HD-shifted spike positions on the HD-shifted trajectory (Fig. 2A second panel). The range of explored shifts was from -6 to +6 cm with steps of 0.2 cm.

#### TIME

In the scenario "TIME" we decreased (past, -) or increased (future, +) the spike times and recomputed spike positions on the recorded trajectory (Fig. 2A third panel). The range of tested shifts was from -600 to 600 ms with 20 ms steps.

#### Distance along trajectory (DAT)

In the DAT scenario we moved spikes by a fixed distance either backwards (-) or forwards (+) along the trajectory. From these new positions on the recorded trajectory we recomputed new spike times (Fig. 2A forth panel). The TIME and the DAT scenarios are related to each other by the animal’s speed. The range of tried shifts was from -6 to 6 cm with 0.2 cm steps.

### Optimization in different dimensions

At every shift we reran the original field detection algorithm. If there was a field detected and if the grid field center was in the 5 cm vicinity of the field center detected at 0 shift, we kept the grid field center and labeled spikes to it as described in ’Grid field detection’. We computed a MI for the field at every shift parameter when there was a detectable field. We chose the optimal shift where MI was closest to 0 (most uniform inbound-outbound tuning).

### Speed division

We divided every cells’ spiking data into 4 equal sized quartiles based on the momentary running speed when spikes were fired. We rerun the field detection algorithm on the 4 quartiles separately and if there was a field center detected in the 5 cm vicinity of a field center of the undivided spiking data we computed the modulation index and the optimal shifts.

### Field size

Field size was computed as:

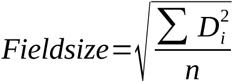

where n is the number of spikes labeled as members of the grid field and D_i_ is the spatial distance of the ’i’th spike position in the field from the grid-field center. All field sizes computed for a grid field during optimization were normalized by the field size computed at 0 shift (FS).

### Experimental design and statistical analysis

We analyzed data recorded by Latuske et al. (2015) and refer the reader to this publication for details on the experimental design. All our analyses were performed in Python 2.7.6. Specific statistical tests used are stated throughout the text. The T-test is taken from scipy.stats. Spearman and Pearson correlations were tested for significance by independently shuffling the two coordinates of the respective data points and computing the corresponding correlation for each new sample. The p-value for correlation is given by the fraction of samples for which the correlation value was larger than in the original sample.

## Results

### Grid-cell data can show non-uniform inbound-outbound tuning

If grid cells encoded the animal’s current allocentric position, a cell’s firing should not depend on whether the animal is running into one of its firing fields, out of the field, or in some other direction relative to the firing-field center. This should even be true for conjunctive cells, which encode location together with the head’s absolute direction in space (Sargolini et al., 2006). On the other hand, deviations from a flat tuning curve in coordinates relative to the grid-field center would indicate that grid cells do not encode the animal’s current position but one shifted in time or space. A similar effect was reported by Huxter et al. (2008), who showed for hippocampal place cells from dorsal CA1 that shifting spike positions along the animal’s current movement direction by about three centimeters leads to the strongest decrease in the firing-field size. In the present study, we wanted to find out whether the shifts needed to obtain flat tuning curves indicate spatial or temporal transformations or even more complicated representations of the animal location.

In the first step, we studied the angular distribution between the animal’s momentary velocity vector and the vector pointing from the spike position to the center of the grid field under study, which we will call the "inbound-outbound angle" (Fig. 1A). Unlike head-direction angles, the inbound-outbound angle represents the orientation of the animal relative to the field center (Fig. 1D) instead of the absolute orientation (Fig. 1C). From these angles we computed an "inbound-outbound firing probability" curve for every grid field, and fitted the data with a cosine model (see Materials and Methods), as shown in Fig. 1E. Based on the modulation amplitude of the cosine fit, we computed modulation indices (MIs) -which measure the modulation amplitude - and peak phases, as described in Materials and Methods. If a grid field’s MI was positive and the peak phase was between -90° and 90°, the field was called "inbound tuned", and "outbound tuned" otherwise (Fig. 1B).

We detected 42 out of 55 significantly tuned grid fields (see Materials and Methods). 40 grid fields had positive MIs and two fields had negative MIs. Most importantly, the mean of the MI distribution (0.46+/-0.27) was highly significantly different from zero (T-test, p = 1.08*10^-13^) as shown in Fig. 1F; the circular mean of the peak phase distribution was -8.55°+/-4.65°, see Fig. 1G. As more than 75% of the grid fields were significantly tuned and more than 95% of the significant fields had a positive MI, there was a clear asymmetry between inbound and outbound firing rates in that grid cells tended to be more active when the animal moved towards the firing field than when it moved away from it.

This phenomenon agrees with the concept of prospective firing (see, e.g., De Almeida et al., 2012).

### Inbound firing rate tuning vanishes when shifting spikes ahead both in space and in time

To explain these observations, we tested four alternative hypotheses:

**MD** ("movement direction"): grid cells anticipate future locations along the momentary velocity vector;

**HD** ("head direction"): the diodes used to track the animal’s location do not represent the position along the head-axis which the grid cells actually encode;

**TIME**: grid cells anticipate future locations using a fixed temporal interval;

**DAT** ("distance along trajectory"): grid cells anticipate future locations along the trajectory with a fixed spatial distance from the current location.

Hypotheses MD and HD assume that a spike is fired in the correct moment, but not for the position where it was recorded. To test whether this could explain the data, we shifted the trajectory points in space and remapped the spikes onto the shifted trajectory (Fig. 2A, first and second subpanel). By altering the size of the shift, we investigated whether approximately flat tuning curves could be obtained. The optimal shifts obtained in that manner might reveal the type of anticipation used and/or the mismatch between the tracing position and the animal’s own sense of self-location.

Similarly, the TIME scenario assumes that a spike was fired earlier or later in time, i.e., prospectively or retrospectively, so we decreased or increased the spike time (Fig. 2A, third subpanel). Finally, the DAT shift assumes that grid cells anticipate along their curved movement trajectory, so we shifted their spikes by a fixed distance along the trajectory (arc length) and recomputed the new spike times (Fig. 2A, forth subpanel). The TIME and DAT shifts are related by the momentary speed.

Every cell’s data were shifted according to the four different scenarios (three when there was no head direction data provided in the recorded data). We shifted both backwards (negative shifts) and forwards (positive shifts). For each shift parameter all the data of one cell were shifted by that given parameter (i.e., if the time shift was -400 ms, all spike times of the cell were decreased by -400 ms).

We only studied optimization curves of grid fields which were significantly inbound-outbound tuned and had a centered peak phase (peak phase less than +/-45° away from 0° or 180°) for zero shift (see also Materials and Methods). We optimized for flat inbound-outbound tuning curves. All four scenarios showed inverted sigmoidal MI optimization curves (Fig. 2B). The optimal shift for a grid field was defined as that shift for which MI was closest to zero, i.e., the cosine fit to the tuning curve at that shift was as flat as possible. Every shifting paradigm resulted mainly in positive optimal shifts, significantly different from zero shift as shown in Fig. 2C (T-test with Bonferroni correction, MD: 3.17+/-1.65 cm, p=3.12*10^-11^; HD: 2.72+/-1.49 cm, p=8.40*10^-8^; TIME: 0.17+/-0.09 s, p=4.03*10^-11^; DAT: 1.97+/-1.95 cm, p=7.78*10^-6^). As illustrated in Fig. 2D, the means of the modulation indices at optimal shifts of all four scenarios were not significantly different from zero (T-test with Bonferroni correction, MD: -0.020+/-0.141, p=0.46; HD: 0.056+/-0.153, p=0.11; TIME: -0.001+/-0.165, p=0.96; DAT: 0.006+/-0.150, p=0.82). We conclude that the optimization indeed flattened the tuning curves such that on average, they are no longer inbound or outbound tuned.

One might expect that the higher modulation index MI is, the higher the expected optimal shift. In other words, when the field is more inbound tuned, flattening it is likely to require a larger forward shift. To test this hypothesis, we calculated the Spearman correlations between the MIs before shifting and the optimal shifts and found that this was indeed true for the MD (r_s_=0.71, p=1.27*10^-5^), TIME (r_s_=0.61, p=3.46*10^-4^) and DAT (r_s_=0.74, p=3.10*10^-6^) shifts as shown in Fig. 2E. For HD we obtained a postive but non-significant correlation (r_s_=0.38, p=0.08). Without the outlier at MI=0.95, the regression line (m=4.7cm, b=1.06 cm, r_s_=0.60) would resemble those for the MD and DAT optimizations.

### Speed-dependency of anticipation

A purely spatial anticipation or a mismatch between the animal’s perceived localization and that measured by the experimenter should not depend on the animal’s speed. However, MI increases, although non-significantly with the mean running speed in the field (r_s_=0.29, p=0.069), suggesting that there could be a temporal aspect in anticipation (Fig. 3A). A closer look at how the optimal shift depends on the animal’s speed shows that the optimal shift for TIME does only change slightly with changing speed (third panel, Fig. 3B). For a rough comparison of the speed dependency in the time-shift paradigm versus those in the other shift-scenarios, the physical dimensions of the slopes, i.e., s^2^/cm versus s, need to be related by a characteristic animal speed, which we take to be 10-15 cm/s (the average of mean movement speed in grid fields is 13.91+/-4.00 cm/s). With this setting, the slope of the regression line of the TIME scenario (m=0.005 s^2^/cm) translates into an effective slope of m=0.04-0.06 s, which is of the same order of magnitude as the MD (0.21 s), HD (0.13 s) and DAT (0.24 s) slopes, but only about a third as large. These rough estimates suggest that the optimal shifts vary less in the TIME scenario than in the other three shift paradigms, with an average temporal anticipation of around 170 ms.

**Figure 3.**
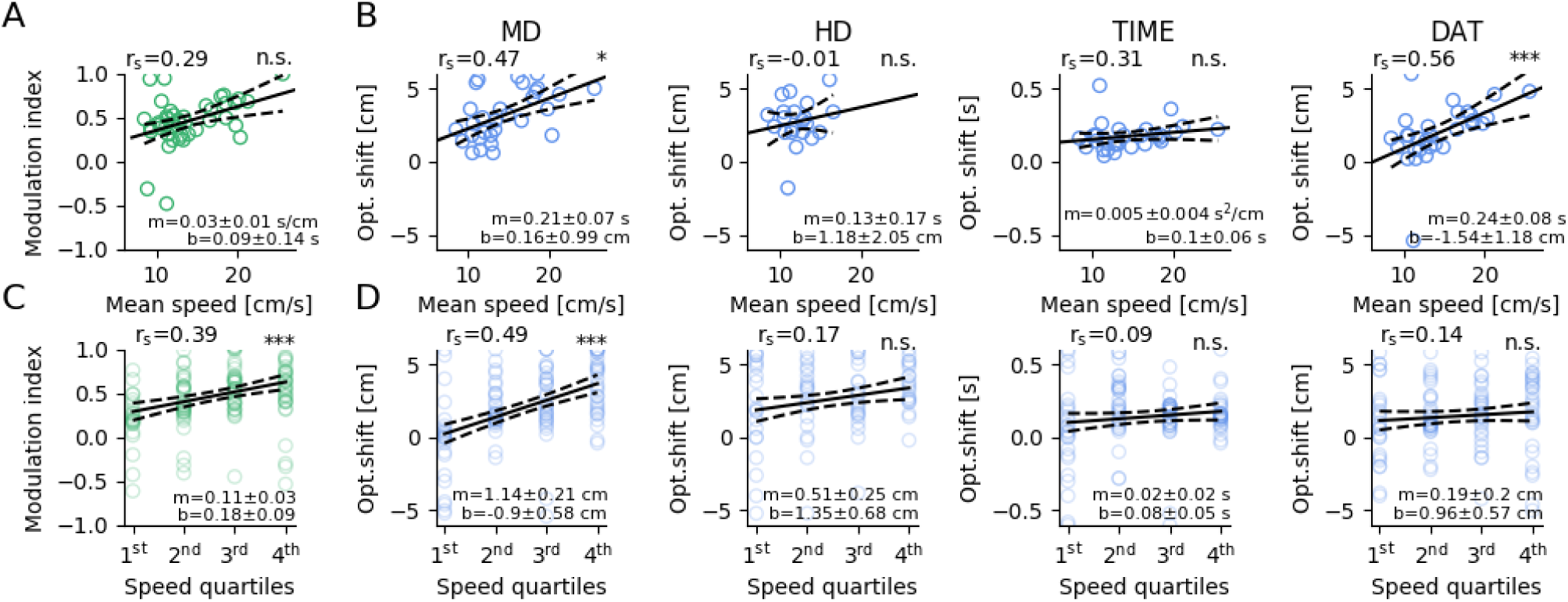
Effect of speed on modulation index and optimal shift A, Modulation indices before optimization as a function of mean running speed within the grid fields. B, Optimal shifts as a function of within-grid-field mean running speed before optimization for the four optimization scenarios. C, Modulation indices of the quartiles before optimization as a function of the quartiles’ rank. D, Optimal shifts of the quartiles as a function of the quartiles’ rank for the four different optimization scenarios. Black solid lines: regression lines, black dashed lines: 95% confidence intervals, r_s_: Spearman correlation values, m: slopes of the regression lines, b: intercepts of the regression lines, single asterisks: significance value for correlation (B: p<0.05/4), three asterisks: significance value for correlation (C: p< 0.01, D: p<0.01/4), n.s.: significance value for correlation (A: p>0.05, B,D: p>0.05/4) (correction for multiple testing).

To investigate whether the speed dependency of the anticipation is true in sessions, we divided every cells’ spiking data into 4 quartiles based on the momentary running speed. We found that indeed, the ranks of the speed quartiles were highly significantly correlated with the corresponding modulation indices (r_s_=0.39, p=6.20*10^-6^) (Fig. 3C). Furthermore, when we correlated the optimal shifts of the quartiles in the case of one spatial scenario, namely MD, we found a highly significant correlation (r_s_=0.49, p=5.19*10^-8^) (Fig 3D). On the other hand, the TIME optimal shifts showed a close to zero, non-significant correlation with the quartiles’ ranks (r_s_=0.09, p=0.34) (Fig. 3D). This suggests that in-session the strength of inbound tuning is speed dependent. This could be explained by a consistent temporal anticipation and thus speed dependent spatial optimal shifts along the trajectory of the animal. More detailed analysis on the level of individual runs through grid fields could not be unambigiously performed, because the spike counts per run would not provide enough spikes for reliably detecting the grid field centers at the tested shift parameters.

### Optimization of firing-field size

Other authors, such as Huxter et al. (2008) and Kropff et al. (2015), have tested grid and place cell coding by shifting spikes ahead either in space or time so as to obtain the smallest grid or place-cell field sizes (FS). We repeated this analysis for the four shifting scenarios (Fig. 4A). Optimal shifts were mostly positive for the smallest field size and the MD, TIME and the DAT optimal shifts were significantly different from zero (T-test with Bonferroni correction, MD: 1.85+/-2.54cm, p=4.79*10-4, HD: 1.65+/-0.86cm, p=0.018, DAT: 1.99+/-2.43 cm, p=1.27*10-4, TIME: 0.115+/-0.226s, p=0.013), as shown in Fig. 4B,C. These shifts are somewhat smaller than those from the inbound-outbound tuning optimization, but the difference if significant only for MD (T-test, MD: p=0.022, HD: p=0.143, TIME: 0.228, DAT: p=0.973). On average, the field sizes decreased to values between 90 and 95% of their starting values (MD: 0.95+/-0.06, HD: 0.91+/-0.06 TIME: 0.91+/-0.08, DAT: 0.92+/-0.06). These results demonstrate that, similar to our results for inbound-outbound tuning curves, firing-field sizes can be optimized in different ways without a particular preferred shift dimension. In other words, being able to reduce the firing-field size along a certain dimension does not imply that this dimension is in any way unique.

**Figure 4.**
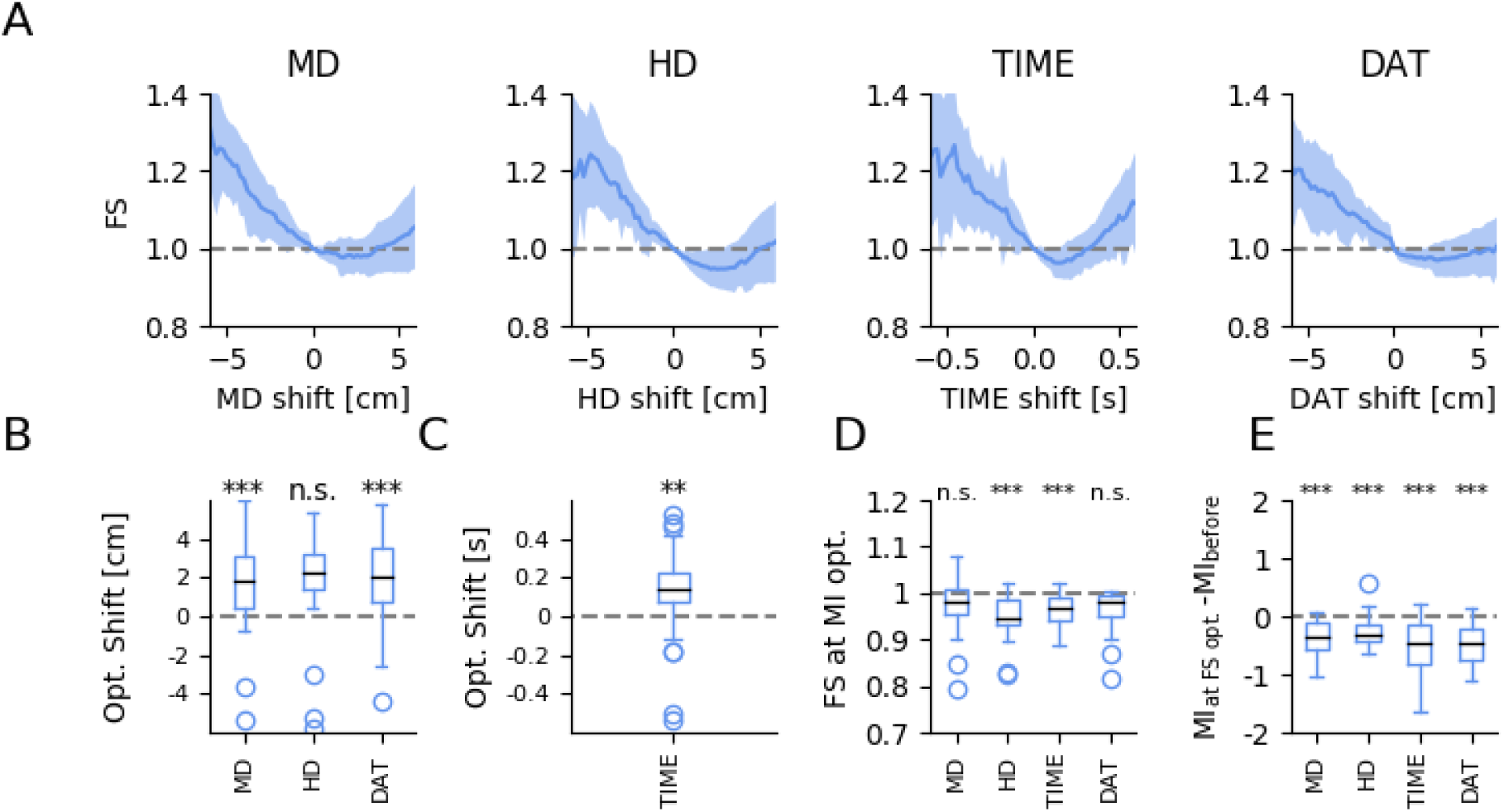
Optimizing firing fields A, Mean optimization curves for the relative field size (FS) in the different scenarios. FS is shown as a function of the tested shift (Blue line: mean MI at shift, shaded area: SD around the mean, Grey dashed line: relative FS at zero shift, i.e., unity). B, Distributions of spatial shifts when optimizing for FS (n.s.: not significant, T-test, p>0.05/3, three asterisks: T-test, p<0.01/3). C, Distribution of optimal TIME shifts for the smallest FS (two asterisks: T-test, p<0.02). D, FS distribution when optimizing for the modulation indices to be close to zero (n.s.: not significant, T-test, p>0.05/4, three asterisks: T-test, p<0.01/4). E, Distribution of the differences between initial MI and optimized values when small field sizes are the target of optimization. (three asterisks: T-test, p<0.01/4)

We then asked how strongly the optimization for MI or FS differed. When we optimized for the modulation index to be close to zero, all four optimization scenarios showed decreased average field sizes that were highly significant for HD and TIME (T-test with Bonferroni correction, MD: 0.98+/-0.06 p=0.037, HD: 0.95+/-0.05 p=1.12*10-4, TIME: 0.97+/-0.03 p=1.64*10-6, DAT: 0.97+/-0.06 p=0.031), see Fig. 4D. When we minimized the FS (Fig. 4E), on the other hand, the average modulation indices, which initially had an average of 0.48 for MD, TIME and DAT, and 0.42 for HD, decreased strongly (T-test, MD: 0.36+/-0.30, p=4.89*10^-7^, HD: 0.25+/-0.26, p=3.29*10^-4^, TIME: 0.51+/-0.47, p=2.45*10^-6^, DAT: 0.48+/-0.34, p=2.69*10^-8^) towards near-zero values (MD: 0.12, HD: 0.18, TIME: -0.02, DAT: 0.01). This finding shows that, indeed, optimizing for FS goes hand in hand with achieving small modulations in the inbound-outbound tuning curves. Thus, although the two optimization schemes involve fundamentally different measures, they nevertheless result in rather similar final configurations.

### Combined optimization along the HD and TIME dimensions

Although a constant temporal shift is consistent with a speed-dependent spatial shift we cannot rule out that both the displacement of the LEDs relative to animal’s perceived location and temporal anticipation lead to the high modulation indices of inbound-outbound tunings. Therefore, we ran a combined tuning-curve optimization where we varied both along the HD and TIME-dimension and picked the optimal combination of spatial and temporal shifts (parameter ranges: TIME: -0.1 s to +2.0 s in 0.02 s steps, HD: -1 cm to +5 cm in 0.2 cm steps). The two-dimensional optimizations resulted in a continuum of optimal shifts (Fig. 5A). This continuum resembles a straight line, in agreement with a linear relation between the HD and TIME components of the joint shift (r=-0.998, p=8*10^-52^). The slope of the linear fit to the average optimization plane (-21.74 +/- 0.23 cm/s) is of the same order though somewhat larger than the typical running speed of the studied mice (population-averaged median speed in grid fields: 13.47+/-4.44 cm/s) which might suggest that the trade-off between a spatial shift and temporal anticipation is related to the animals’ movement characteristics. The continuum of optimal solutions underscores the difficulty in assigning a unique cause to the inbound-outbound tuning of grid cells.

**Figure 5.**
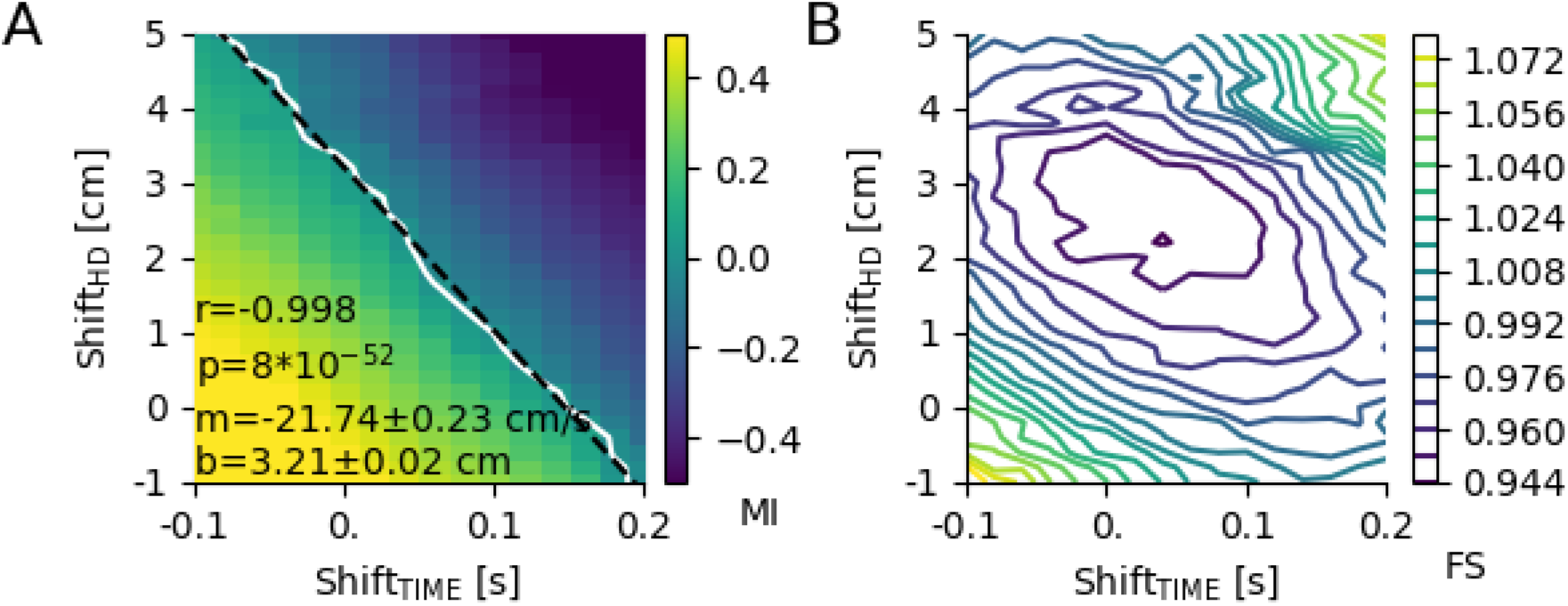
Population average of combined HD and TIME optimizations A, When spikes are simultaneously shifted along the HD and TIME dimension, a continuum of optimal solutions with vanishing MI emerges, here shown for the average across all analyzed grid fields. This continuum illustrates the trade-off between spatial mismatch (LED position vs. perceived location) and temporal anticipation. White line: zero-crossing of the mean optimization plane, black dashed line: regression line, r_s_: Pearson correlation value, p: p-value for Pearson correlation, m: slope of the regression line, b: intercept of the regression line B, When joint shifts along the HD and TIME dimension are used to minimize firing-field size, a shallow minimum can be observed (HD shift: 2.79+/-1.72 cm, TIME shift: 0.024+/-0.070 s).

When one simultaneously minimizes the firing-field size along the HD and TIME dimensions, a shallow minimum appears at the population level (Fig. 5B). From a parabolic fit of the data for FS<1, optimal spatial shifts were calculated as 2.31 cm, optimal temporal shifts were 0.036 s. The two-dimensional HD-TIME optimization does not lead to significantly better improvements in FS compared to the one-dimensional HD or TIME optimizations (p=0.87 and p=0.85, respectively). The ellipsoidal near-optimal field-size contours in Fig. 5B are tilted and again suggest a trade-off between temporal anticipation and spatial offset, in qualitative agreement with the tuning-curve results.

## Discussion

Analyzing the dataset of Latuske et al. (2015) we found that grid cells in mice fire prospectively (positive MI) almost exclusively. This result differs from findings by De Almeida et al. (2012) who reported that rat grid cells can also be in a retrospective mode. Furthermore, the spatial anticipation measured by De Almeida et al. was up to 15 cm, whereas pure spatial shifts calculated in this paper are around 3 cm. The differences might partly reflect differences in animal size as well as differences in the experimental design – fast straight runs on linear tracks without foraging in the study of De Almeida et al. versus curved trajectories during foraging in an open arena in the study of Latuske et al. – that result in different behavioral states.

To better understand the inbound-outbound firing characteristics in the dataset of Latuske et al. we studied four different scenarios – anticipation along the movement direction (MD), head direction (HD), TIME and DAT (distance along trajectory). The optimal spatial shifts measured in the MD, HD, and DAT paradigm were around 3 cm. Shifts in that range were also reported by Huxter et al. (2008) who recorded place cells in rat hippocampus (CA1) and minimized the place-field size along the movement direction. These authors then assumed that the apparent prospective firing results from a mismatch between the LED position and the animal’s perceived location. However, minimizing firing-field sizes can again be done along the same dimensions as minimizing tuning-curve modulations (Fig. 4). For all four dimensions investigated, shifting spikes can reduce the firing-field size – as for tuning-curves, there is no special role for shifts along the movement direction. This finding implies that the place-cell data of Huxter et al. (2008) might possibly also have an interpretation as true temporal anticipation or some mixture of spatial and temporal components, as in our work. The temporal anticipation scenario was also supported by our finding that in-session the modulation indices and the MD spatial optimal shifts are speed dependent. This could be explained by the consistent TIME optimal shifts which were shown to be speed independent (Fig. 3).

For rat CA1 place-cell ensemble coding a theta-sequence look-ahead with a goal-dependent extent was reported by Wilkenheiser and Redish (2015). Since in the experimental data analyzed in our study there were no goals present the results cannot be directly compared. However, the prospective fashion of the coding is supported by those findings. For grid cells in the superficial layers of medial entorhinal cortex of rats, indications of a similar look-ahead have been found as well (O’Neill et al. 2017). In addition, recent evidence suggests that hippocampal theta sequences extend to MEC and represent forward-directed sweeps as well as left-right alternations of prospective trajectories in egocentric coordinates (Gardner et al., 2019). The detailed relation between these phenomena and the modulated inbound-outbound tuning curves studied here has not been addressed yet.

The optimal TIME shift found in the present study is around 170 ms, which clearly differs from the 70 ms reported by Kropff et al. (2015) for rat grid cells with theta-modulated speed cell input (T-test, p=1.29*10^-6^). In both studies, however, the spatial anticipation along the trajectory showed the same qualitative running-speed dependency. Further analysis is needed to reconcile the quantitative differences. On the other hand, when we optimized for minimal field size, as in Kropff et al. (2015), the distribution of temporal shifts was not significantly different from 70 ms (T-test, p=0.31). In other words, optimizing for small field size results in smaller optimal shifts than when focusing on inbound-outbound tuning.

The present exploratory study underscores the importance of distinguishing between spatial aspects and temporal anticipation and points to a trade-off once both types of shifted activity are jointly considered for optimizing the inbound-outbound tuning curves. Given the optimization criteria (i.e. the modulation index) should be the closest to zero and not reaching a local extremum, the two dimensional inverted sigmoidal optimization plane has many zero crossings. This trade-off could be especially relevant when the animal is taking straight runs through a grid field. In these cases the temporal and the spatial forward shifts would be both along the same directions. In such scenarios, the spatial and temporal anticipation are hard to distinguish. Furthermore, the trade-off would be more obvious if the running speed of the animal during these runs is relatively constant. In addition, comparison with the published literature shows that there remain a number of open issues. We do hope that our investigation will trigger follow-up studies to clarify these questions.

## Author contributions

All authors designed research. DÉC performed research and analyzed data. DÉC and AVMH wrote and edited the paper with support from JN and MS.

